# A prize-collecting Steiner tree application for signature selection to stratify diffuse large B-cell lymphoma subtypes

**DOI:** 10.1101/272294

**Authors:** Murodzhon Akhmedov, Luca Galbusera, Roberto Montemanni, Francesco Bertoni, Ivo Kwee

## Abstract

**Background:** With the explosion of high-throughput data available in biology, the bottleneck is shifted to effective data interpretation. By taking advantage of the available data, it is possible to identify the biomarkers and signatures to distinguish subtypes of a specific cancer in the context of clinical trials. This requires sophisticated methods to retrieve the information out of the data, and various algorithms have been recently devised.

**Results:** Here, we applied the prize-collecting Steiner tree (PCST) approach to obtain a gene expression signature for the classification of diffuse large B-cell lymphoma (DLBCL). The PCST is a network-based approach to capture new insights about genomic data by incorporating an interaction network landscape. Moreover, we adopted the ElasticNet incorporating PCA as a classification method. We used seven public gene expression profiling datasets (three for training, and four for testing) available in the literature, and obtained 10 genes as signature. We tested these genes by employing ElasticNet, and compared the performance with the DAC algorithm as current golden standard. The performance of the PCST signature with ElasticNet outperformed the DAC in distinguishing the subtypes. In addition, the gene expression signature was able to accurately stratify DLBCL patients on survival data.

**Conclusions:** We developed a network-based optimization technique that performs unbiased signature selection by integrating genomic data with biological networks. Our classifier trained with the obtained signature outperformed the state-of-the-art method in subtype distinction and survival data stratification in DLBCL. The proposed method is a general approach that can be applied on other classification problems.

## BACKGROUND

Diffuse large B-cell lymphoma (DLBCL) is the most common form of human lymphoma. Based on gene expression profiling studies, it has been proposed that this clinically heterogeneous disease can be divided in at least two main classes, the germinal center B-cell (GCB) and the activated B-cell like (ABC), subtypes, depending on their phenotypic homologies with their supposed cell of origin (COO) [36, 12]. These subgroups have shown differences in clinical course and response to standard therapies, therefore, they should allow the development of tailored therapies [36, 12]. The definition of the COO is now recommended in the clinical practice [33] although it still a matter of research on the identification of the best markers to be used and on the choice of the best technology to measure them [36, 21, 15]. Recent advances in genome sequencing instruments have now provided researchers with a huge amount of data from which it is possible to extract useful gene expression signatures to discriminate ABC and GCB subtypes of DLBCL. Nevertheless, the analysis of gene expression patterns is characterized by a large number of covariates, a large number of genes compared to a small number of observations, and a limited number of patients with known ABC or GCB subtypes. Therefore, data mining must be accomplished with careful statistical techniques to identify a signature that contains only the elements most useful for the subtype prediction.

There are several signature sets proposed in the literature to identify the subtypes of lymphoma [9, 16, 7, 31, 26, 28, 10], but the lists of genes in these sets are very discordant. We demonstrate the gene overlap between four signatures in **Fig. 1**, where the numbers in the figure indicate the amount of overlapping genes. These signatures are strongly dependent on the specific study and sometimes determine the discordant COO definition of the same sample. One main reason for this heterogeneity could be the preliminary filtering process of genes when a subset of all genes analyzed is selected to reduce the degrees of freedom that allows further statistical analysis. We propose a PCST-based method capable of extracting important genes by searching the complete gene space from the expression or transcriptomics datasets, thereby removing the initial arbitrary gene filtering step. Basically, the PCST approach enables the analysis of genomics data at a network level, which leads to the identification of subnetworks that may possibly act as a gene expression signature to distinguish the DLBCL subtypes. We construct gene interaction networks using gene co-expression data without applying any gene filtering as a whole, and score them using the differential expression [13]. Afterwards, the PCST is applied to find a neighborhood of genes in the network that are differently expressed and connected to each other through the highly mutual informative (∼ reliable) edges. The set of differently expressed genes that are collected with high mutual information among themselves would potentially stratify the DLBCL patients.

**Figure 1:**
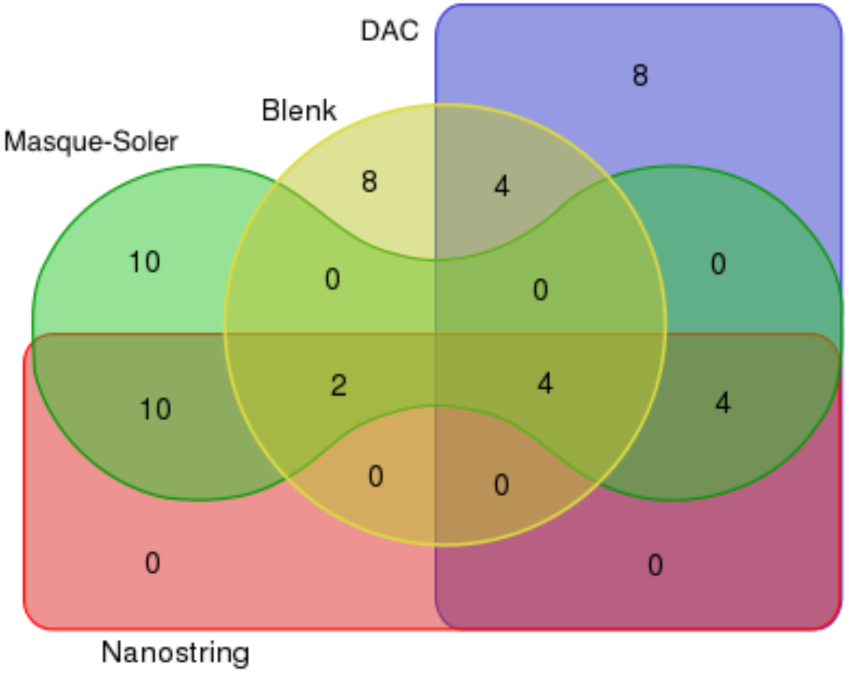
A Venn diagram demonstrating the gene overlap from the published signatures including (DAC [9], Blenk [7], Masque-Soler [26], and Nanostring [30]). Numbers in the figure indicate the amount of overlapping genes between the signatures. There is a considerable discordance in the gene lists. The diagram is made using the online tool (source: www.bioinformatics.psb.ugent.be).

## METHODS

### Datasets

We used seven publicly available datasets downloaded from the Gene Expression Omnibus^1^ (GEO) repository that are reported in Table 1. The first three columns of the table contain the names, the generated platforms and the number of samples in the datasets. The datasets have been generated in different experiments where the source is provided in the last column of the table. As reported in the fourth column, the first three datasets are used to obtain the PCST signature and to train the regression-based model, ElasticNet [47], to perform the classification. The rest of the datasets are used to compare the performance of the ElasticNet model trained with the PCST signature against the golden-standard approaches from the literature. For detailed information about the datasets used to obtain the signature and to train the regression model, see Table 2.

**Table 1:** Publicly available datasets. The first three columns of the table contain the names, generated platforms and number of samples in the datasets. The datasets have been generated in different experiments where the source is provided in the last column of the table. We also specify under the fourth column which dataset are used to identify the PCST signature and to train the ElasticNet regression model [47], and later on to test the performance of the ElasticNet trained with the PCST signature.

| Dataset | Platform | Samples | Use | Author |
| --- | --- | --- | --- | --- |
| GSE22470 | Hgu133A | 176 | Train | [29] |
| GSE19246 | Hgu133Plus2 | 144 | Train | [43] |
| GSE31312 | Hgu133Plus2 | 451 | Train | [14] |
| GSE4475 | Hgu133A | 96 | Test | [17] |
| GSE10172 | Hgu133A | 10 | Test | [20] |
| GSE23501 | Hgu133Plus2 | 57 | Test | [32] |
| GSE10846 | Hgu133Plus2 | 350 | Test | [22] |

**Table 2:** The datasets that are used to obtain the PCST signature and train the ElasticNet [47]. The total number of samples in each data set includingthe number of ABC and GCB subtype samples is provided in the table.

| Dataset | Total samples | ABC samples | GCB samples |
| --- | --- | --- | --- |
| GSE19246 | 144 | 63 | 81 |
| GSE22470 | 215 | 71 | 144 |
| GSE31312 | 451 | 214 | 237 |

To prepare the data sets for subsequent analysis, we transformed the probe names into hugo gene symbols [27]. The probe is a complementary sequence that targets the certain location of a genome while hugo gene symbols contain a unique naming of the specific locations of a genome [27]. If more than one probe is associated to the same gene, only the probe with the highest standard deviation is retained. To utilize the genes with some missing values, we imputed missing values for the genes lacking measurement in less than 50% of the samples using a nearest neighbor average method [38]. Other genes with more than 50% missing values across the samples are removed from the computations, because imputing missing values for these genes could introduce a considerable amount of artifacts.

### Networks

To obtain the PCST signature, we generated two interaction networks for each of the three training datasets using the multiplicative model of the ARACNE [25] algorithm. A total of six interaction networks are generated with up to 21’049 nodes and 1’423’267 edges as reported in Table 3. ARACNE [25] is a powerful tool for the reconstruction of gene interaction networks. It applies the data processing inequality to all triplets of nodes in the network to remove the least significant edge in each triplet using the threshold parameter *τ*. For each triplet of nodes *i, j* and *k*, the weakest edge *e*_*ij*_, is removed from the network if its weight is smaller than *e*_*ik*_ *τ* and *e*_*jk*_ *τ*. Since we wanted to analyze the generated networks in a broader context, we selected a less stringent threshold for *τ*. We generated two interaction networks for each training data with parameters *τ* = 0.01 and *τ* = 0.05. In these networks, each edge represents the interaction between two genes and ARACNE provides a score for each edge based on the mutual information of corresponding gene expression values. The node prizes are labeled with the differential expression for each gene as *p*_*v*_ = |*E*_*ABC*_ – *E*_*GCB*_|, where *E*_*ABC*_ and *E*_*GCB*_ are equivalent to the mean of gene expressions for ABC and GCB cancer patients, respectively. The edge costs are set to *c*_*ij*_ = 1- *m*_*ij*_/*m*_*max*_, where *m*_*ij*_ represents the mutual information between the expression values of gene pairs *i* and *j*, and *m*_*max*_ is the maximum mutual information within the datasets.

**Table 3:** The sizes of network instances generated by the training datasets. The networks are generated by the multiplicative model of ARACNE algorithm [25] using the parameters values of *τ* = 0.01 and *τ* = 0.05.

| Dataset | $eps = 0.001$ | | $eps = 0.005$ | |
| --- | --- | --- | --- | --- |
|  | Nodes | Edges | Nodes | Edges |
| GSE19246 | 20639 | 82282 | 20639 | 154469 |
| GSE22470 | 12994 | 266085 | 12994 | 375097 |
| GSE31312 | 20639 | 1055055 | 20639 | 1423267 |

### PCST - the network optimization method

The PCST is a well-known problem in graph theory and there are different variants of the problem in the literature [1, 2, 23]. Given an undirected graph *G* = (*V, E*), where the vertices are labeled with prizes *p*_*v*_ ≥ 0 and the edges are labeled with cost *c*_*e*_ *>* 0, the goal is to identify a subnetwork *G*′= (*V*′, *E*′) with the tree structure. The target is to minimize the sum of the prizes of vertices out of *V*′, and to minimize the total cost of the edges in *E*′. This is equivalent to the minimization of the following objective function [19]:

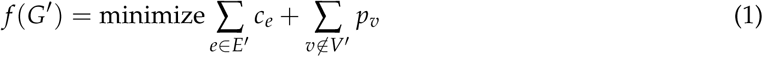

The PCST has a wide range of applications, mainly in the design of utility networks such as fiber optic and district heating networks. Recently, some PCST applications in biological networks have been published [4, 39, 40, 41]. The biological graphs such as genetic interaction networks can be scored by using experimental data including gene expression profiling, mutation, or copy number data. In these networks, every vertex is a gene, and every edge between two vertices represents the genetic or protein interaction between the two vertices. Each vertex in the network is given a prize, which can be the differential expression between two subtypes or number of mutation for that gene [13]. Also, each edge in the network is given a cost, which corresponds to pairwise mutual information of expression values between two genes. After having scored all vertices and edges in the network, the PCST is employed to detect a relevant subnetwork (tree). Biologically, the subnetwork obtained by the PCST has an important meaning [4], in which it corresponds to a portion of interaction network where many genes are closely related in terms of their functions, and potentially belong to the same biological pathways [13]. Furthermore, the resulting subnetwork contains many genes that are highly correlated and differently expressed between the sub-types in the experimental data. Therefore, the set of genes included into PCST solution tree can represent gene expression signatures to distinguish disease subtypes.

The illustration of the PCST on input graph and the resulting output tree are demonstrated in **Fig. 2**. The vertex sizes and edge widths are proportional to prize of nodes and mutual information of edges, respectively.

**Figure 2:**
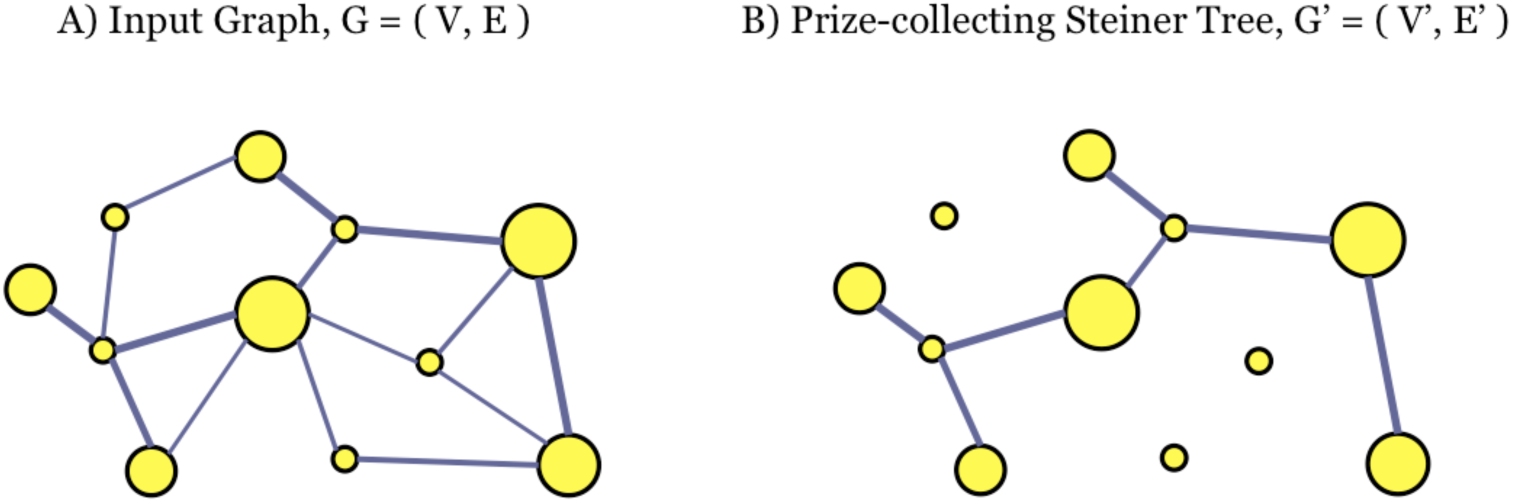
The Prize-collecting Steiner Tree problem instance and solution. A) An input graph instance. The node sizes and arc widths are proportional to the prizes and mutual information, respectively. B) The Prize-collecting Steiner Tree solution for input graph. The PCST collected set of genes that have higher differential expression and connected through the least cost (∼highly mutual informative) edges.

In the application of PCST in genomics, it is possible to have very large networks to analyze. Nevertheless, PCST has NP-hard characteristics which makes it computationally challenging for existing exact algorithms to find solutions in reasonable execution time. This yields a need for fast matheuristic algorithms to compute the PCST solution tree in large networks for inferring biologically relevant sub-networks. Considering challenges, previously we have developed a matheuristic approach [1, 2] which efficiently scales up on large biological instances. The matheuristic was composed of the preprocessing procedures, a heuristic clustering algorithm and an exact PCST solver from the literature [23]. Here, we employ a prior approach developed in [1] to compute the PCST solution tree for biological networks.

### Signature

The PCST method described in [1] is applied in six interaction networks to obtain solutions. Basically, a PCST solution includes itself the group of genes that are differently expressed in ABC versus GCB subtype, and have high mutual information among themselves at the same time. Therefore, these genes are assumed to be a promising gene expression signature to stratify ABC and GCB subtypes. Furthermore, we combine the PCST results of multiple datasets by selecting the intersecting genes in order to get a more robust signature.

### Training the classification methods

We use ElasticNet [47] as a classification method to distinguish the DLBCL subtypes. The classification method is trained by using the signature obtained from the PCST as described in Section, and its performance is examined on the test data to objectively evaluate it. We use the default parameter values in ElasticNet.

#### Penalized linear regression and mixture of experts

The idea is to look for a simple linear classifier performing a logistic regression. The Steiner tree allowed us to reduce the number of covariates from tens of thousands of genes to ten, however among them there could still be correlated genes or genes not relevant for the classification, which can lead to a sub-optimal estimation of the regression parameters. To select relevant genes several techniques of variable selection and penalized linear regressions have been devised to identify the important covariates. We used ElasticNet regression [47] that combines a continuous variable selection, while keeping under control the correlations among the covariates. The penalization introduced by ElasticNet can be seen in a Bayesian framework as the introduction of a prior distribution on the values of the coefficients which models our prior assumption that the value of the coefficients must be bounded (this keeps under control any correlation among the regressors) and that the solution could be sparse (variable selection). This penalization is an attempt to combine the well-known Lasso and Ridge methods, trying to overcome their individual limitations.

Moreover, we adopted a meta-analytic approach in which each training dataset is used to build an independent classifier. Each classifier performs a binary classification through a logistic regression and for every sample it casts a vote between 0 and 1 that can be interpreted as the probability for the sample to be of sub-type ABC. We tried two different ways to combine the votes of the single classifiers together. Either we used the mean value or the median; we found that the mean leads to best results (see Results and Supplementary Material).

Another question is how to present the training data to the classifiers. In fact, PCA might allow to reduce the dimensionality of the datasets and improve the prediction power of the ElasticNet model. As it can be seen in **Fig. 3**, if we consider the three training datasets restricted to the genes present in the signature found, the first three components are enough to explain more than 60% of the total variance, so we chose to consider only the first three components.

**Figure 3:**
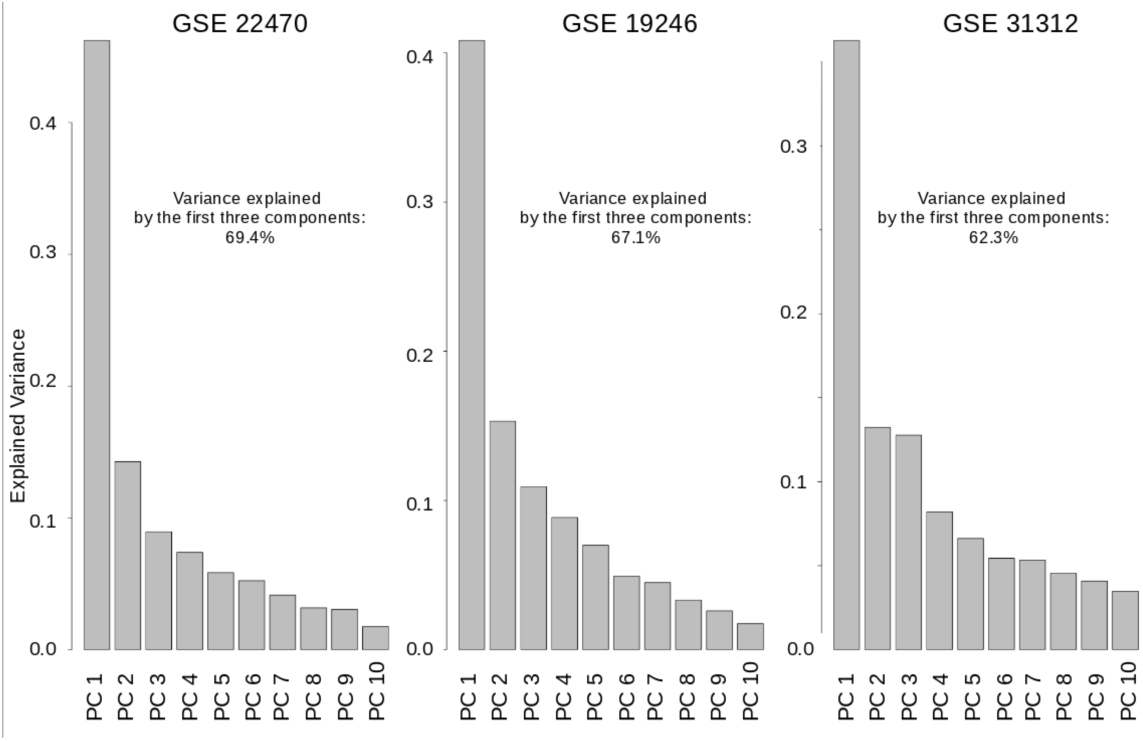
The PCA of the three training datasets shows that the first three components are enough to explain more than 60% of the total variance.

We tried to feed the classifier using either the real data or the first two principal components and we found that the best results are obtained using both representations (See Results and Supplementary Material for a more detailed discussion).

In the end the classification algorithm works as follows:

1. For each training dataset, train an ElasticNet classifier;
2. For each training dataset, train second ElasticNet classifier based on the first three principal components;
3. Let each classifier cast a vote for each sample in the test dataset;
4. Merge the votes together using the mean value.

### Comparison with previous studies

We compared our gene expression signature and classifier with previously proposed ones. The comparison was on two phases: comparison of the classification accuracy and patient survival stratification ability.

To test whether our signature and classification algorithm can improve the present classification system, we tested it against the DLBCL Automatic Classifier (DAC) [9] one of the “state-of-the-art” classifiers for the DLBCL classification.

Since ABC and GCB DLBCL groups have different clinical outcome we compared the stratification using our classes and using the classes assigned by the original study, for two of the test datasets with available survival information.

## RESULTS and DISCUSSION

We compared the performance of our gene expression signature with previous methods in the literature. The performance comparison was in twofold: i) comparing the classification accuracy of DLBCL patients provided by the methods and ii) comparing the survival stratification ability of the methods. In the first comparison, we provide accuracy results of the methods by benchmarking our method against the DLBCL Automatic Classifier (DAC) [9]. Currently, the DAC is the “state-of-the-art method” in the literature to classify DLBCL patents. Alternatively, in the second comparison, we report the patient stratification power of our method based on survival data.

### Comparison to previous signatures

As discussed in the introduction, various signatures have been proposed to classify ABC and GCB subtypes of DLBCL in the literature. These signatures barely overlap, however, we figured out that they are highly correlated in terms of biological processes in which they are involved. With the R package 

~~~
nclust
~~~

 [45, 44], we performed a hierarchical clustering [24] of genes for all datasets using the distance metric 1 – *cor*, where *cor* indicates the Pearson’s correlation [6] between the genes. We then computed the fold-changes for each gene, which is the mean difference in expression of ABC and GCB subtypes. We ordered the fold-changes according to the dendrogram provided by the hierarchical clustering, resulting in a signal exhibiting some peaks that were smoothed with a Gaussian kernel [3], which corresponds to highly correlated and differentially expressed genes between the two subtypes. Using this ordered fold-change signal as the backbone, we plotted the previous signature sets from the literature in **Fig. 4**, which provides a unified overview of different signatures. In the figure, the signature genes are distributed under the peaks of the ordered fold-changes. Although the signatures do not contain overlapping genes, they have several genes belonging to the same peaks, i.e. very correlated. This supports the hypothesis that even if the signatures are different, they contain genes belonging to the same functional group that can represent the same biological function.

**Figure 4:**
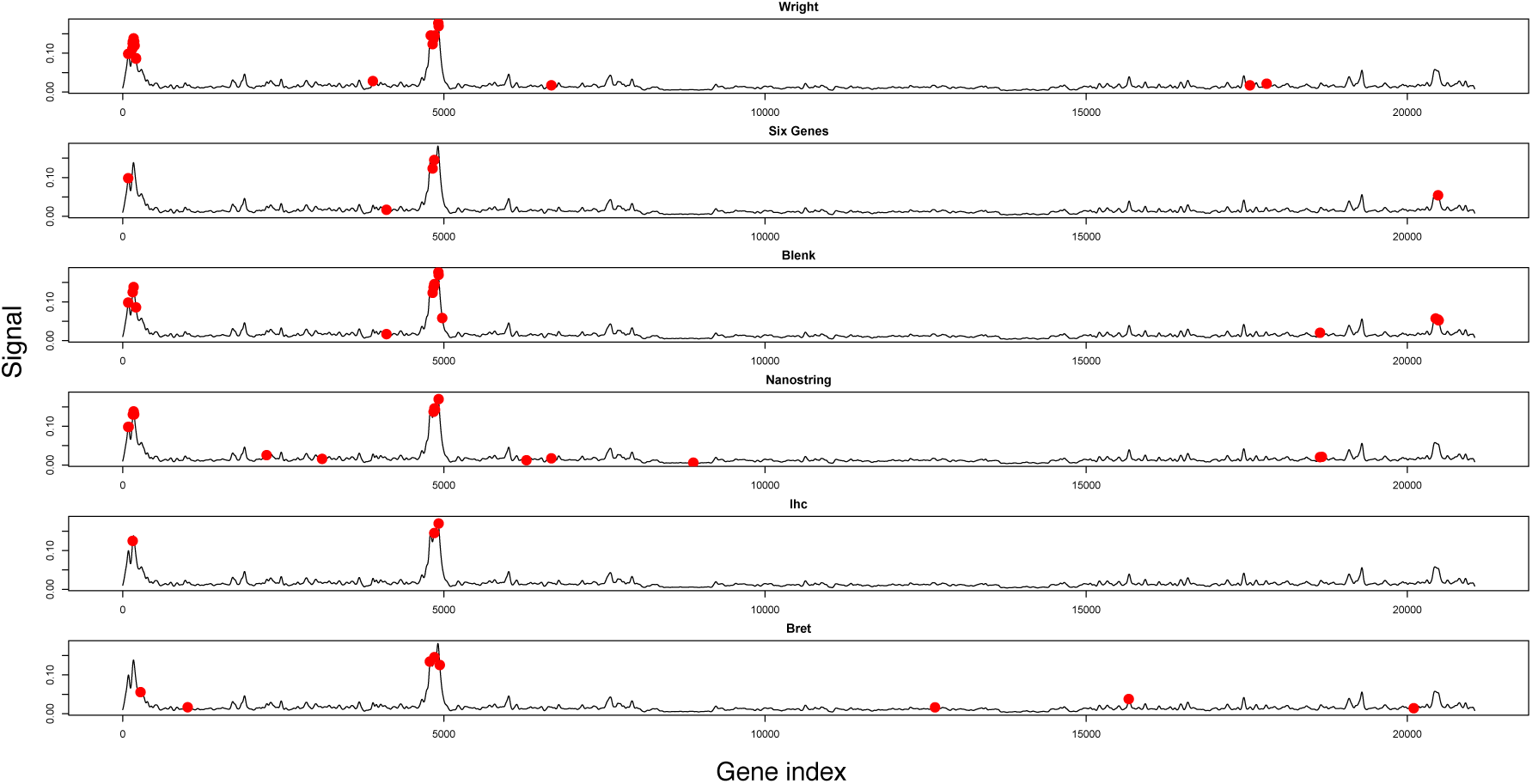
Signatures including Wright [46], Six Genes [18], Blenk [7], Bret [8], IHC [16] and Nanostring are displayed in the figure based on the ordered fold changes according to the the hierarchical clustering dendrogram.

The immunohistochemical (IHC) [16] analysis is the golden standard for early diagnostic purposes of DLBCL patients, while the DAC classifier [9] implements the Wright [46] signature genes to classify ABC and GCB subtypes.

### The PCST signature

For illustrative purposes, the union of PCST subnetworks for six interaction networks is displayed in **Fig. 5**, resulting in a network of 174 genes and 240 edges. In order to obtain a robust signature set, we selected intersecting genes in six PCST subnetworks, including ten genes, namely: *MYBL1, PHF16, SSBP2, SPINK2, RAB7L1, STAG3, C13ORF18, BATF, MME* and *LMO2*.

**Figure 5:**
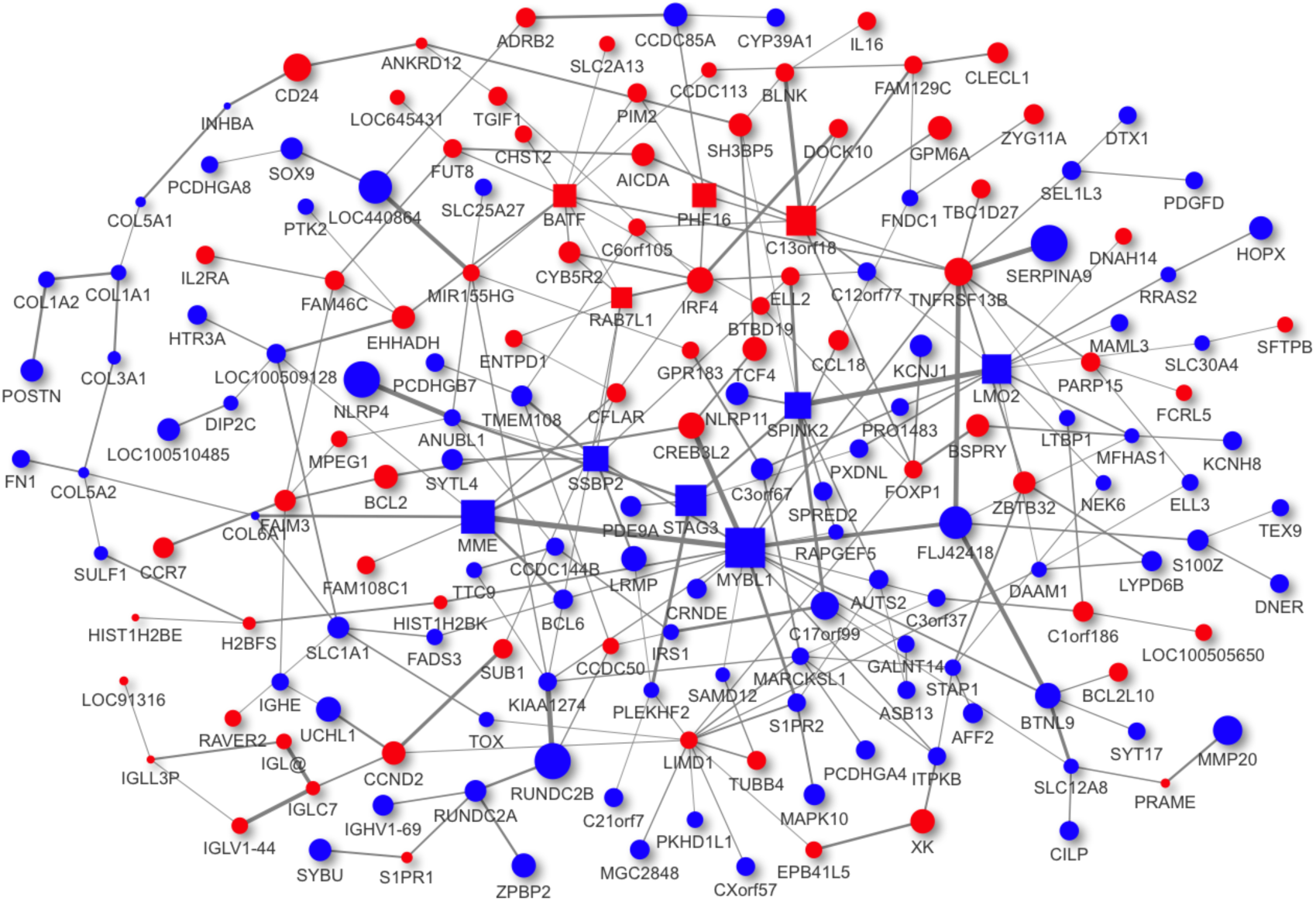
The subnetworks obtained by the PCST are overlaid on top of one another. The node sizes and edge widths are proportional to the differential expression of genes and mutual information between the genes, respectively. Red genes are over-expressed in ABC, and blue genes are over-expressed in GCB subtype. The square-shaped genes are selected as the PCST signature.

**Fig. 6** shows the overlap of our PCST signature genes with the two main peaks of the ordered fold-changes signal, which is described in the previous section.

**Figure 6:**
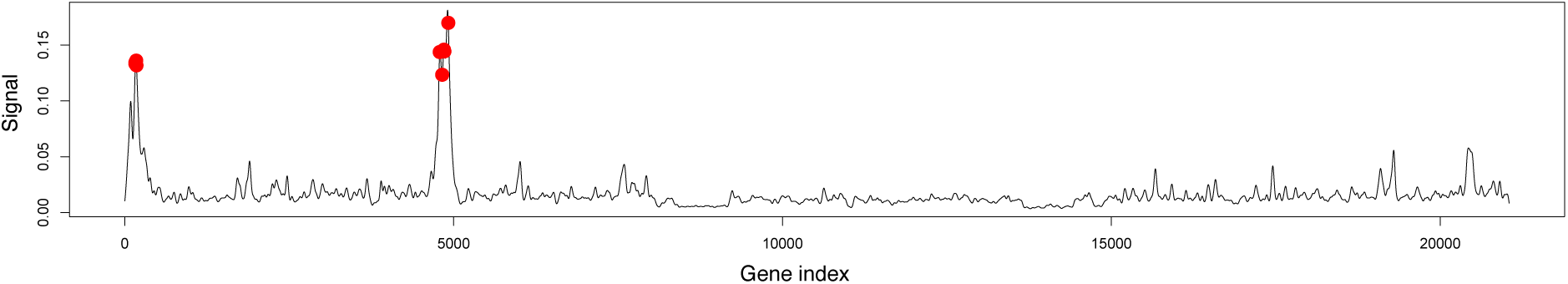
Our signature genes coincide the two main peaks of the ordered fold-change signal. These main peaks are also shared by the other signatures found in the literature (see also **Fig. 4**).

Furthermore, we compared the expression levels of genes belonging to the PCST signature between ABC and GCB subtypes in **Fig. 7**. The signature genes were differentially expressed between the subtypes throughout the training datasets, demonstrating the potential distinctive power of the signature in ABC and GCB subtypes.

**Figure 7:**
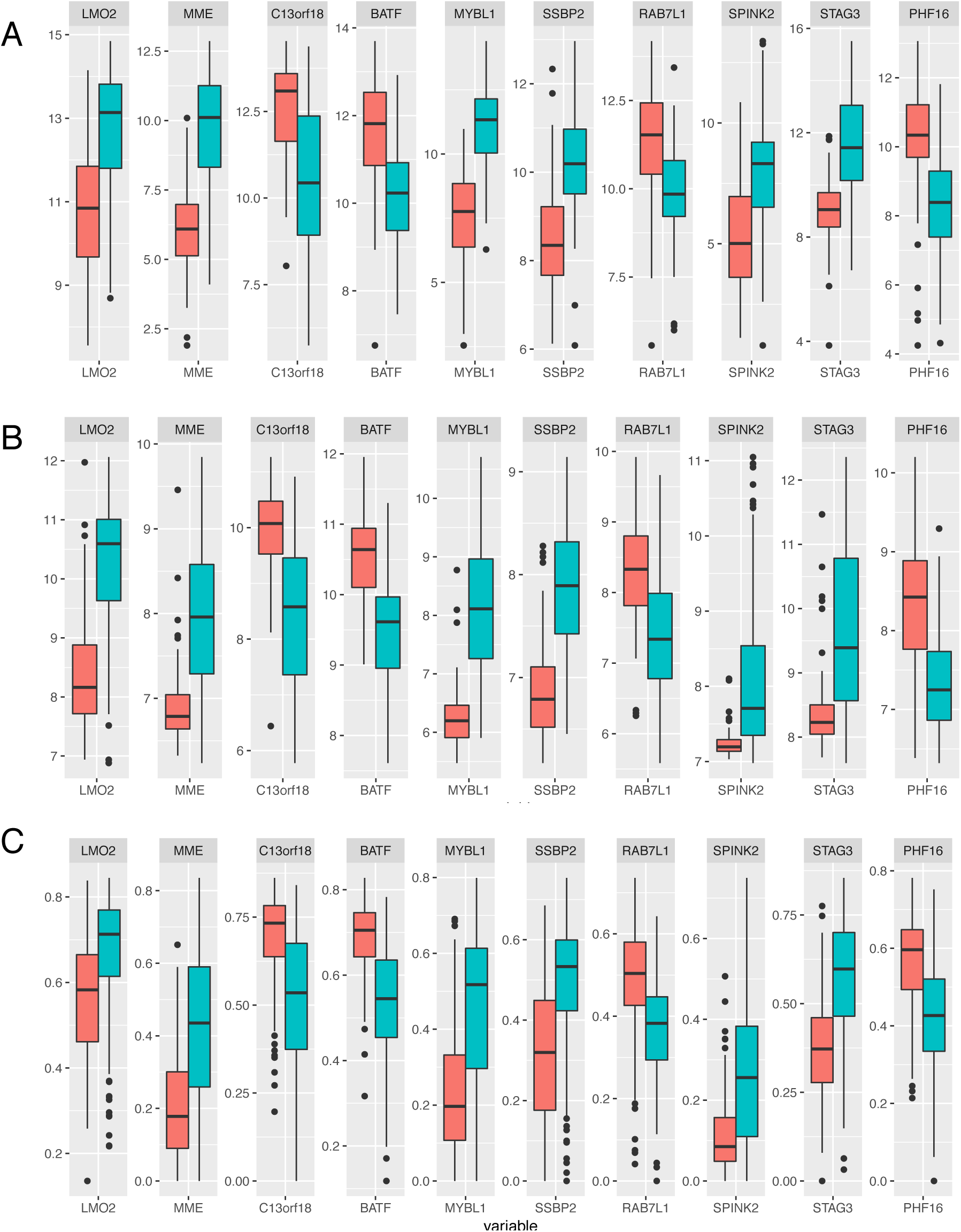
The mean values of ABC (orange) versus GCB (turquoise) subtype expressions for the PCST signature genes A) for the GSE19246 dataset, B) for the GSE22470 dataset and C) for the GSE31312 dataset.

### Accuracy comparison with previous models

We used the ElasticNet regression model [47] trained with our signature to classify DLBCL patients and compared its performance with the DAC [9] classifier. The accuracy is computed by dividing the number of correctly predicted patients by the total number of patients. Other minor subtypes of DLBCL than ABC and GCB are considered as the type-III subtype within the DAC classifier. To create a fair comparison platform, we disregarded the type-III subtype predictions of the DAC while computing the accuracy, or reassigned the type-III subtype to the ABC or GCB based on the highest prediction probability.

The comparison results for four test datasets are reported in Table 4. Both methods provided similar accuracy results for the first two datasets in the table. However, our method significantly outperformed the accuracy of the DAC classifier [9] for the last two datasets. We used the procedure described in Methods, but in the Supplementary material section we showed the results of our classifier trained with different settings.

**Table 4:**
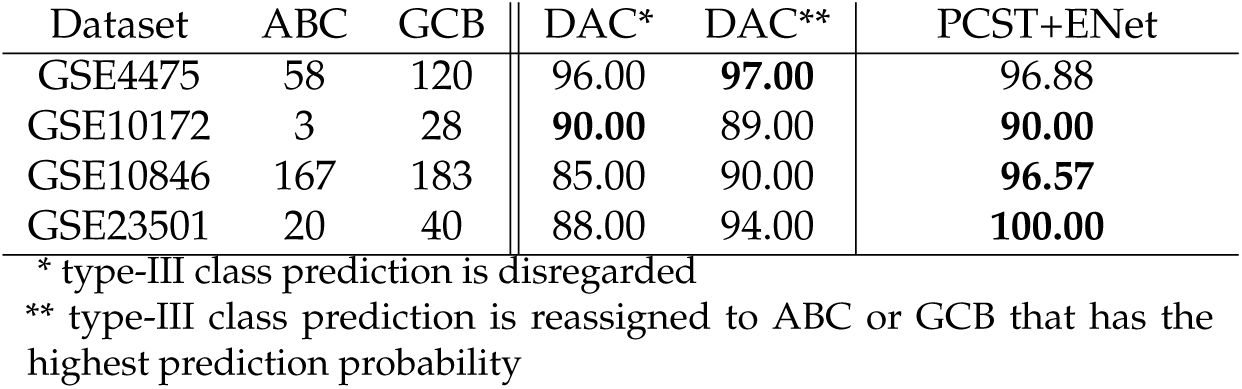
Survival analysis for two datasets. The top and bottom rows contain the results for GSE4475 and GSE10846 datasets, respectively. The left column shows the survival analysis using the classes provided by [17] and [22] while the right column demonstrates the performance of our PCST signature classification.

### Survival analysis with the PCST signature

To provide a stronger evidence of the PCST signature relevance in clinical trials, we tested the signature in the analysis of survival data. The signature was used to stratify DLBCL patients using survival data. Two out of four datasets within the training datasets contained the survival data in the GEO repository. The survival analyses are displayed in **Fig. 8** for these datasets, where our method provided better p-values (1.14e-9 versus 2.56e-9 [22] for GSE10846 and 0.0000984 versus 0.00296 [17] for GSE4475) in the stratification of DLBCL patients compared to previous approaches.

**Figure 8:**
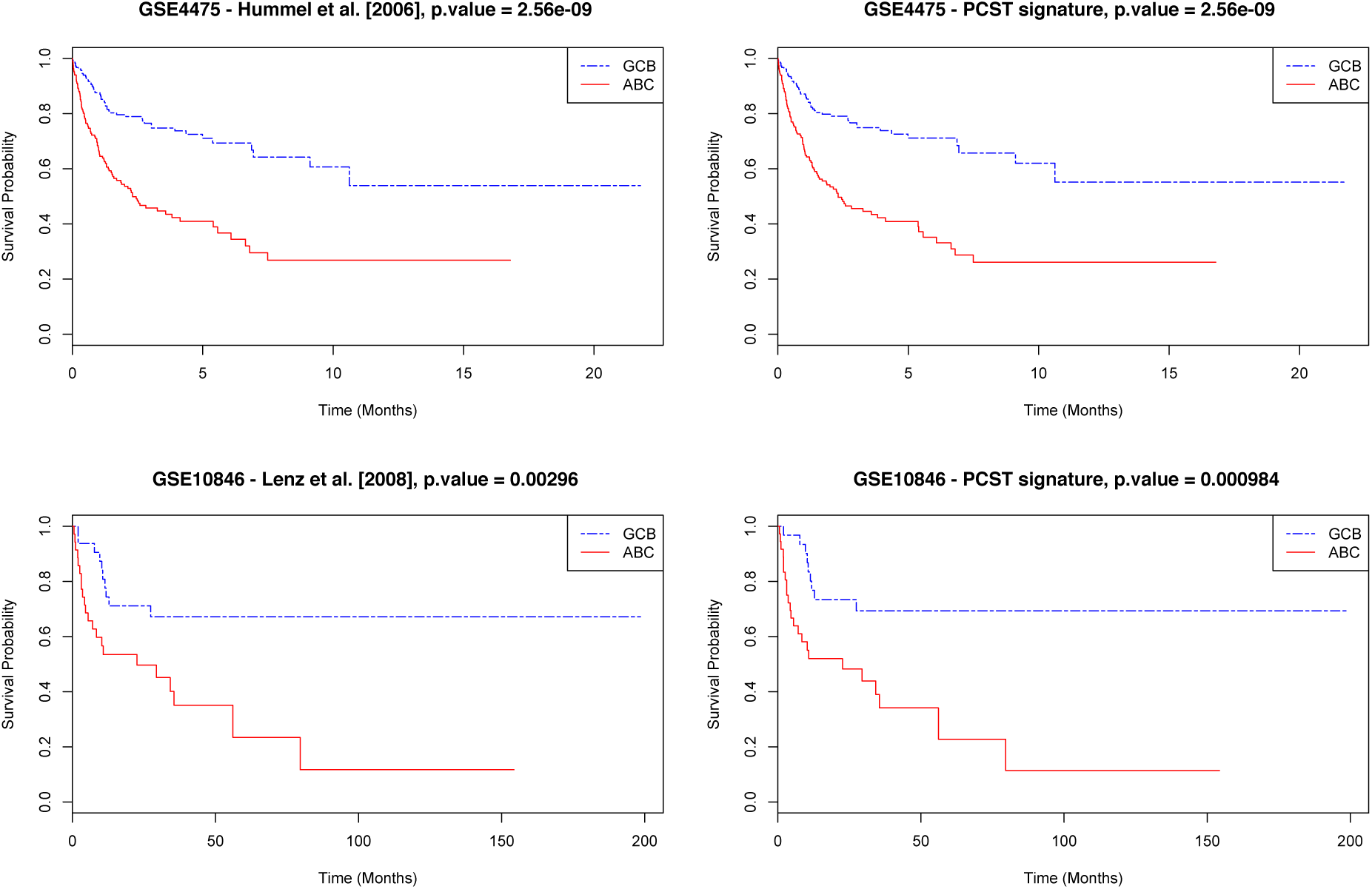
Survival analysis for two datasets. The top and bottom rows contain the results for GSE4475 and GSE10846 datasets, respectively. The left column shows the survival analysis using the classes provided by [17] and [22] while the right column demonstrates the performance of our PCST signature classification.

## Conclusion

High-throughput genomics is a promising technology that enables personalized medicine. However, the way to make it available for usual clinical practice is still far from being completed. One main challenge in high-throughput data analysis is to identify signature genes that can stratify patients with relevant clinical outcomes. Here, we proposed a method based on the well-known Steiner Tree algorithm. The genes analyzed by the specific platform are mapped onto the vertices of a genetic network where the vertex prize is given by the differential expression of the gene between the two subtypes of disease (hence it is a measure of how good the gene is able to discriminate the two subtypes) and the edges are given a cost corresponding to the mutual information between two genes connected by the vertex. The aim of the PCST is to find a relevant subnetwork with minimal cost; this subnetwork is comprised of genes with high discriminative power and mutual information. Since the problem is NP-hard, we employed a metaheuristic previously developed in our group [1, 2].

We identified the gene expression signature for the DLBCL patients with our method and tested it against the “state-of-the-art” approach in the literature. The classification accuracy of the proposed method was equivalent or better than the “gold-standard” method. In addition, our method resulted in better stratification of DLBCL patients using the survival data.

Finally, the proposed feature selection method is robust to noise in data due to the batch effects and other environmental factors. It is very generic method that can be easily applied to other types of cancer data for signature selection and subtype stratification.

## Supplementary

### Publicly available signatures

In the literature, various signatures have been proposed in order to classify the DLBCL. In **Fig. 4** we showed how the genes from these signatures overlap with the peaks of the hierarchical clustering. The signatures used are

1. Wright: One of the first signatures [46].
2. Six Genes [18]: Minimal signature composed of only six genes
3. Blenk [7]: Signature of 18 genes
4. Nanostring [30]: Signature used to classify FFPE tissues using the Nanostring technology.
5. IHC [16]: Signature for IHC classification
6. Bret [8]: 12 genes signature

### Using different parameters to train the classifier

In the main test we showed the classification accuracy using a mixture of experts composed on models trained on the real data and on the first three principal components. The final vote is given by the mean of the single experts.

However, one can wonder if using both real data and the first three principal components is not redundant, or if the median is better than the mean since it is less dependent on outliers. We present in Table 5 the results using either the mean or the median on classifiers trained only with real data, only with the three first principal components or both. As it can be seen, the mean values of classifiers trained both on real data and on the principal components allows to achieve the best performance. We also report the results obtained with the DAC [9] classifier.

**Table 5:**
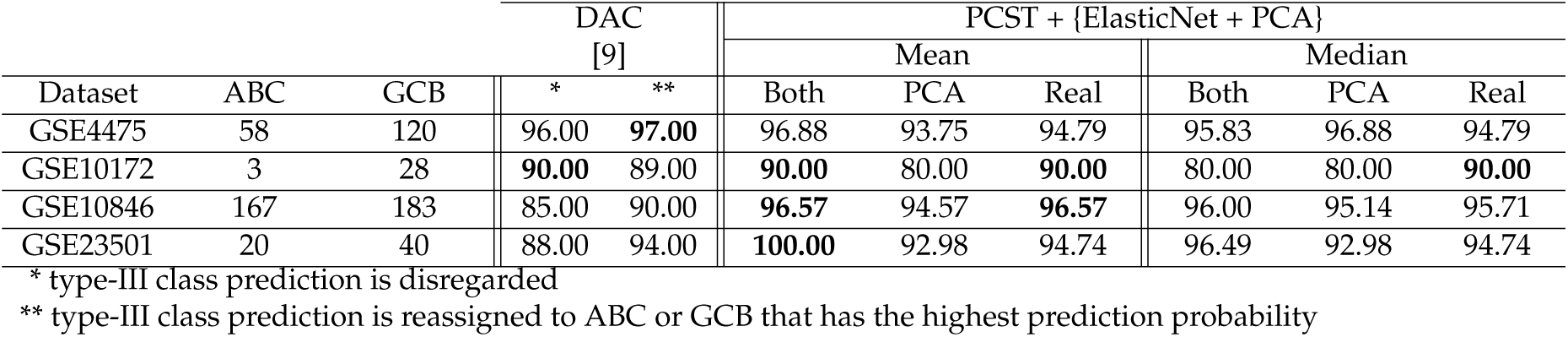
Comparison of the mixture of experts using ElasticNet. The different mixtures depend on how the training data are feed into the classifiers and how the final vote is obtained as a combination of the votes of the single classifiers.

## Abbreviations

PCST: Prize-collecting Steiner Tree
GEO: Gene Expression Omnibus
DLBCL: Diffuse large B-cell lymphoma
ABC: Activated B-cell like
GCB: Germinal center B-cell
COO: Cell of origin
DAC: DLBCL Automatic Classifier

## Acknowledgements

Not applicable

## Availability of data and materials

All the gene expression data used during the current study is publicly available on Gene Expression Omnibus. The software for developed signature selection and classification is available from the corresponding author on request.

## Author’s contributions

M.A. and L.G. designed the experiments, performed all analysis and wrote the paper. M.A., I.K. and R.M. developed the PCST algorithm. I.K. and F.B. conceived the experiments and provided the biological interpretation.

## Consent for publication

Not applicable

## Ethics approval and consent to participate

Not applicable

## Footnotes

1 http://www.ncbi.nlm.nih.gov/geo/

